## Supplementary information for "Abasy Atlas v2.2: The most comprehensive and up-to-date inventory of meta-curated, historical, bacterial regulatory networks, their completeness and system-level characterization"

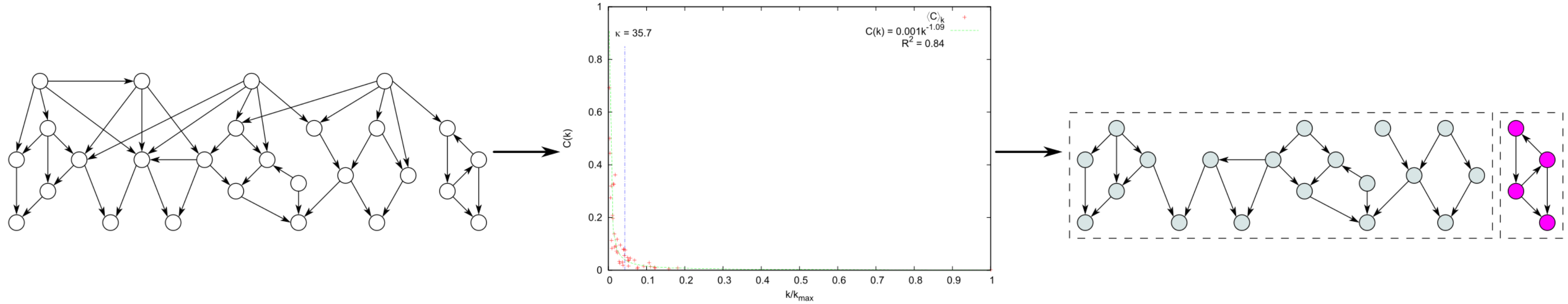


A regulatory network is represented as a directed graph. Each gene is a node, while arcs between them stand for regulatory interactions.

Global regulators are identified by the κ-value, which defines an equilibrium point between two contradictory behaviors: hubness (high out-degree, low clustering) and modularity (low out-degree, high clustering). First, the clustering coefficient distribution, *C*(*k*), is computed. Then, given $C(k) = \gamma k^{-\alpha}$, the κ-value is calculated by using the formula $\kappa = \sqrt[\alpha+1]{\alpha\gamma} \cdot k_{\max}$.

Identification of functional modules. Global regulators (connectivity > κ-value) and their links are removed from the network, thus naturally revealing the modules (isolated islands) and the basal machinery genes (single isolated nodes not encoding for regulators, not shown here for the sake of clarity).

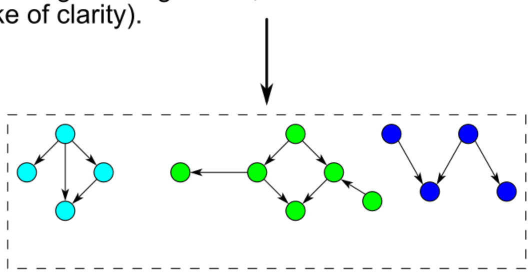

Identification of pre-submodules composing the megamodule. The megamodule is isolated and all the non-regulator-encoding genes and their links are removed, disaggregating it into isolated islands. Each island is identified as a pre-submodule.

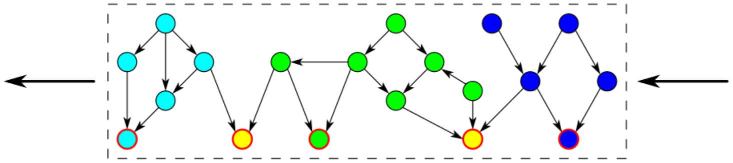

Identification of intermodular genes. All the removed non-regulators-encoding genes (red-outlined nodes) and their interactions are added back following a rule: if gene G is regulated by genes belonging to two or more submodules, then G is classified as intermodular (yellow nodes), else G is added to the same submodule than its regulators.

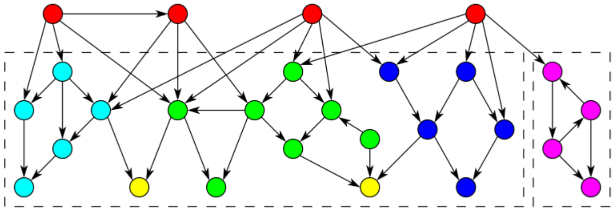

Finally, global regulators (red nodes) and their links are added back, thus reconstructing the original network but additionally revealing its hidden diamond-shaped three-layered architecture and its systems-level components: global regulators, modules, basal machinery and intermodular genes.

**Supplementary Figure 1.** Methodology of the natural decomposition approach (NDA).

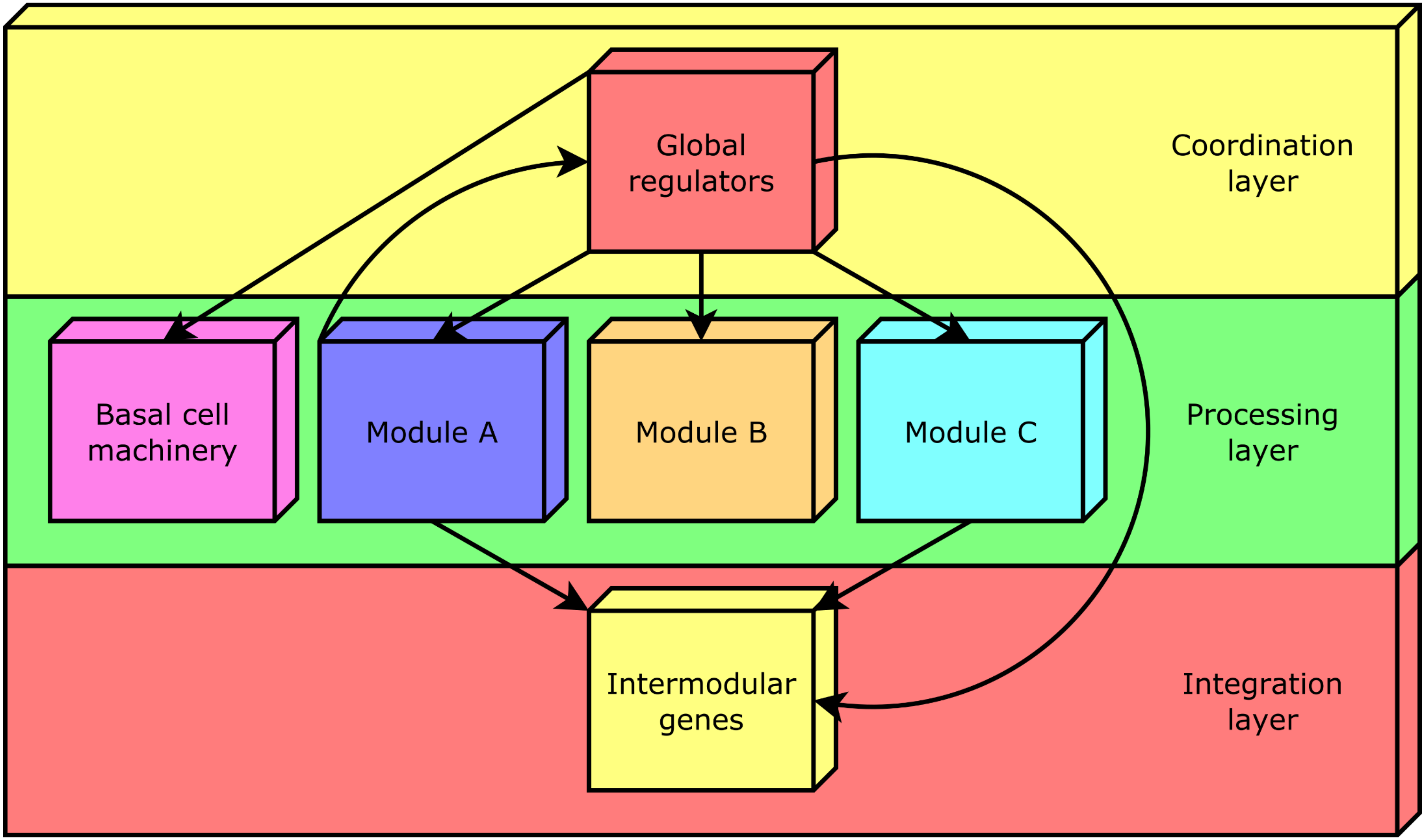


**Supplementary Figure 2.** The functional architecture unveiled by the NDA is a diamond-shaped, three-tier hierarchy, exhibiting some feedback between processing and coordination layers, which is shaped by four classes of system-level elements: global regulators, locally autonomous modules, basal machinery, and intermodular genes.

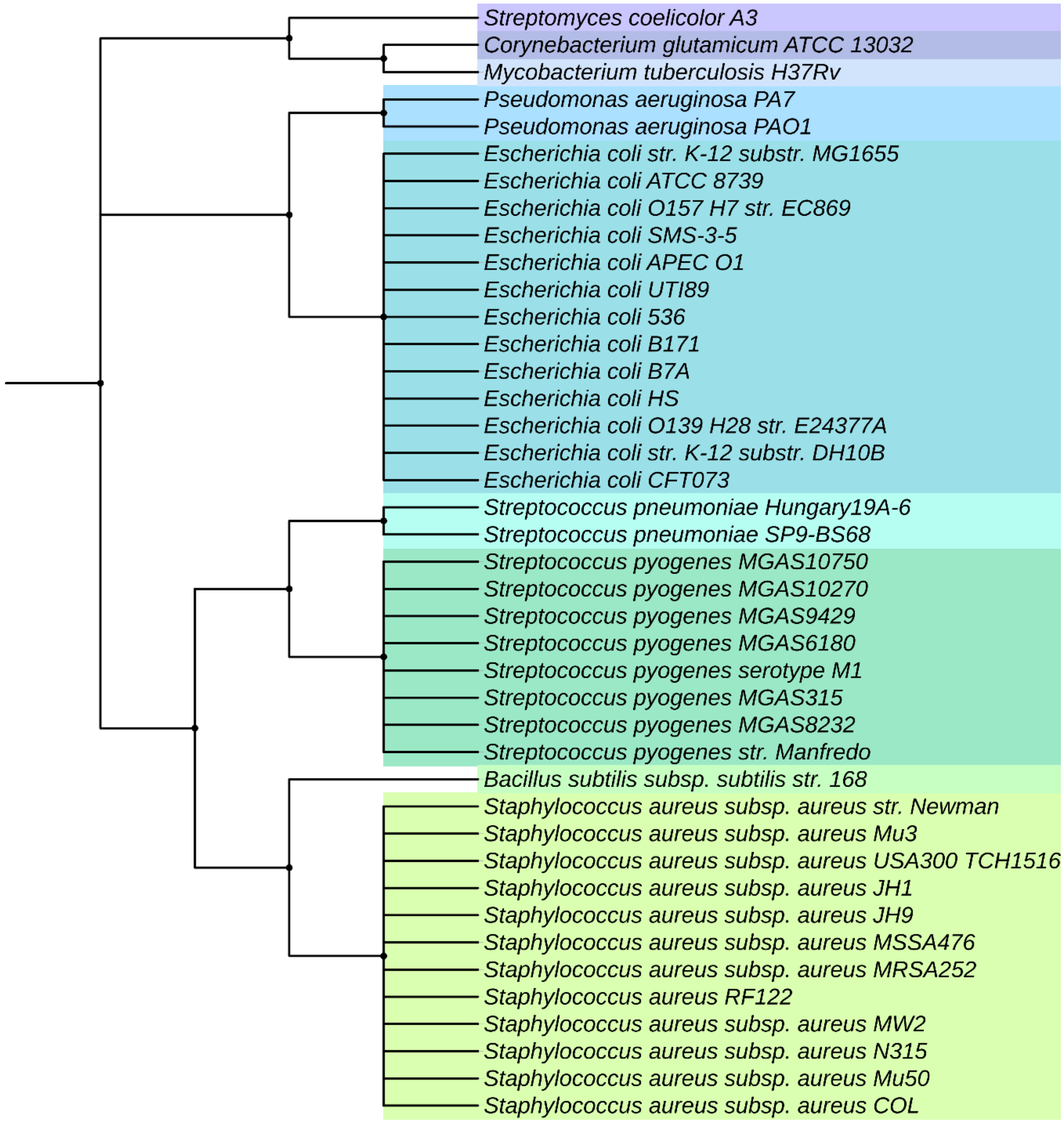


**Supplementary Figure 3.** Abasy integrates gene regulatory interactions for 9 species and 41 strains including model organisms.

| DB abbreviation | PMID | Reference |
|---|---|---|
| **Bsub15** | 26577401 | [1] |
| **CRN12** | 22080556 | [2] |
| **Cglu17** | 27829123 | [3] |
| **DBTBS08** | 17962296 | [4] |
| **Mtuber11** | 21818301 | [5] |
| **Mtuber12** | 22737072 | [6] |
| **Mtuber15** | 25581030 | [7] |
| **Mtuber16** | 27029515 | [8] |
| **Paeru11** | 22587778 | [9] |
| **RDB01** | 11125053 | [10] |
| **RDB04** | 14681419 | [11] |
| **RDB06** | 16381895 | [12] |
| **RDB11** | 21051347 | [13] |
| **RDB13** | 23203884 | [14] |
| **RDB16** | 26527724 | [15] |
| **RDB18** | 30395280 | [16] |
| **RTB13** | 23547897 | [17] |
| **SW16** | 26433225 | [18] |
| **SW18** | 29788229 | [19] |

**Supplementary Table 1.** Name abbreviation and year used as code for the source field in the network ID, and the PMID of the article.

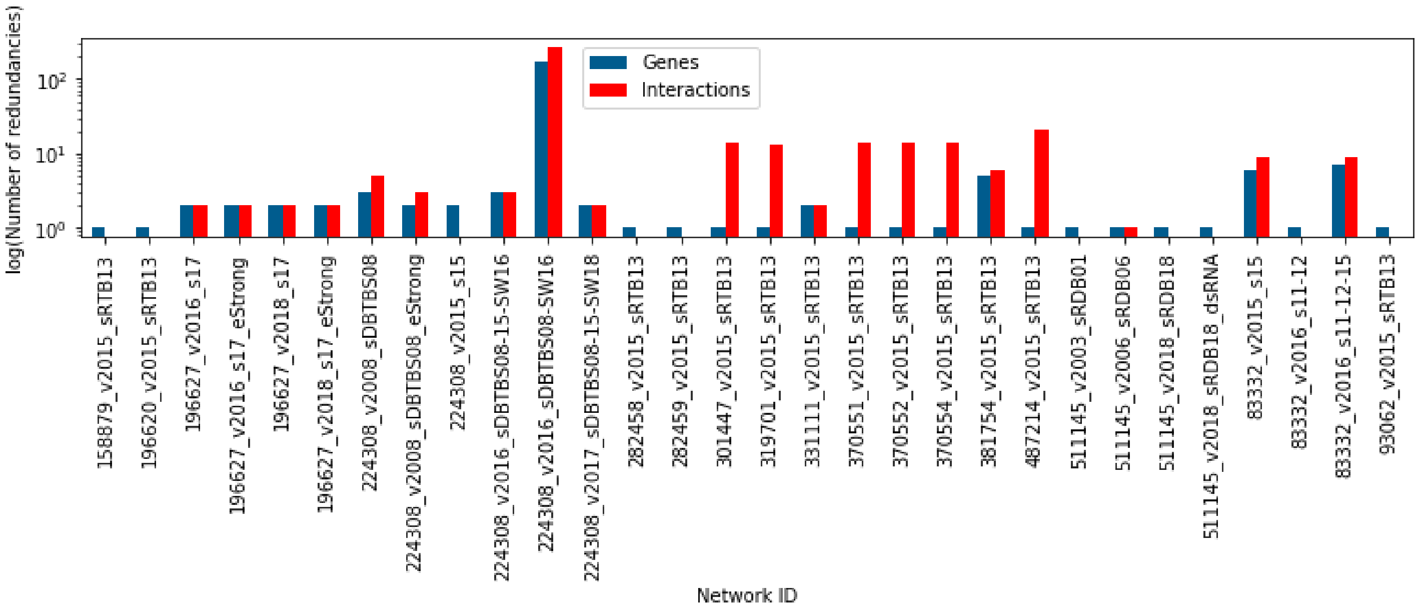


**Supplementary Figure 4.** Number of redundant genes and interactions removed due to the synonyms resolution in the meta-curated GRNs. A total of 223 nodes and 412 interactions are removed.

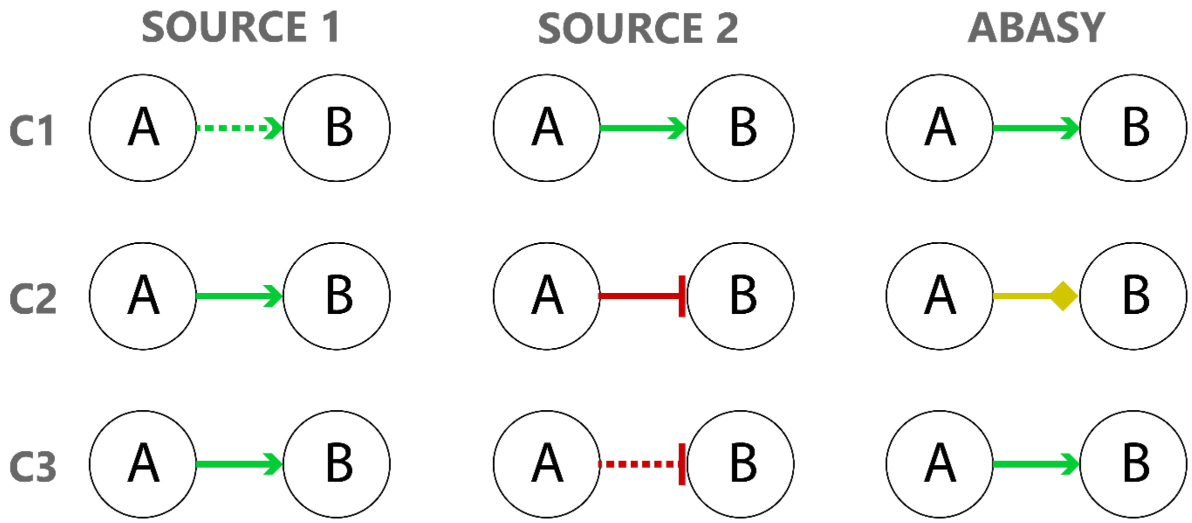


**Supplementary Figure 5.** Information preserved when consolidating regulatory interactions from different sources. **C1:** In case of same effect but different evidence level, the interaction shared and the "strong" evidence is conserved. **C2:** In case of different effects and the same evidence level, both effects are conserved in a single dual interaction to avoid redundancy. **C3:** In case of both attributes are different, only the "strong" interaction is conserved.

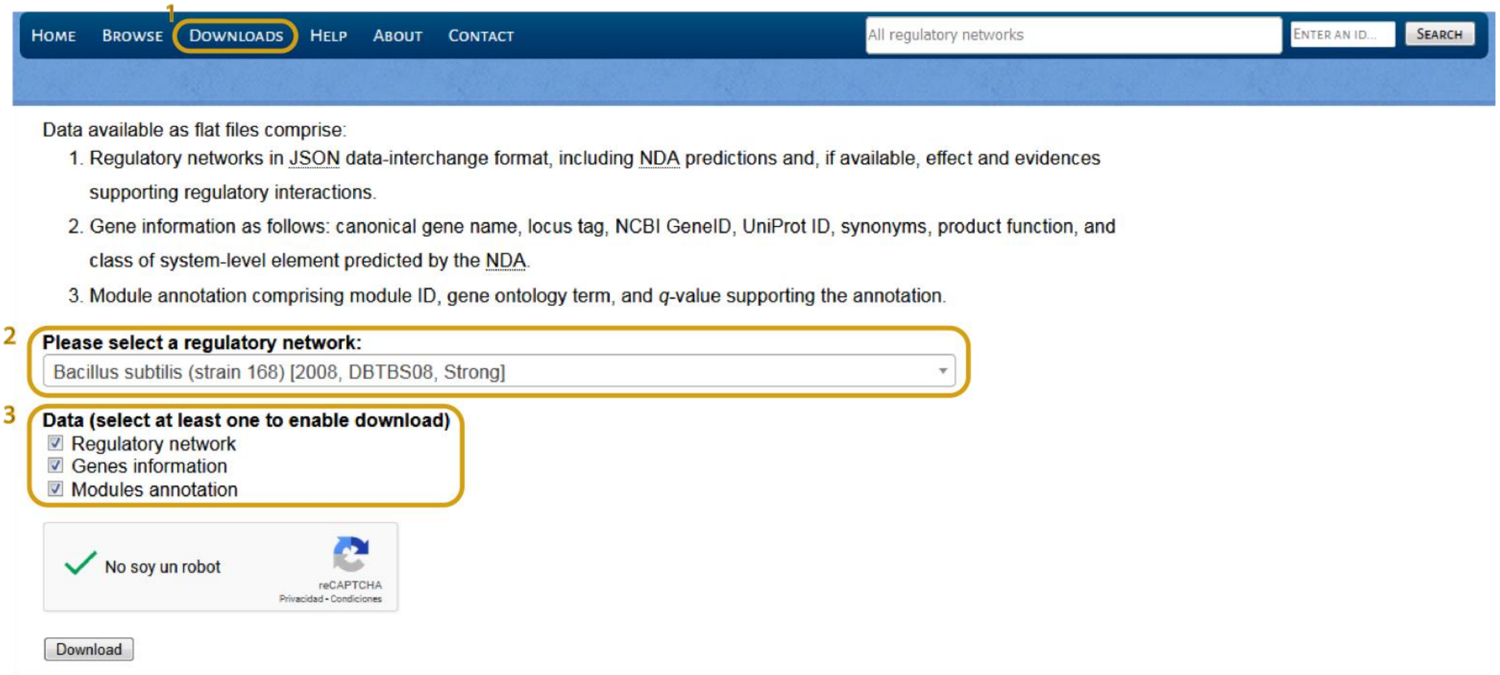


**Supplementary Figure 6.** Filtered GRNs to contain only "strong" interactions van be also downloaded through the "Downloads" page where you can select the GRN you want as well as the additional data such as gene information ad modules annotation.

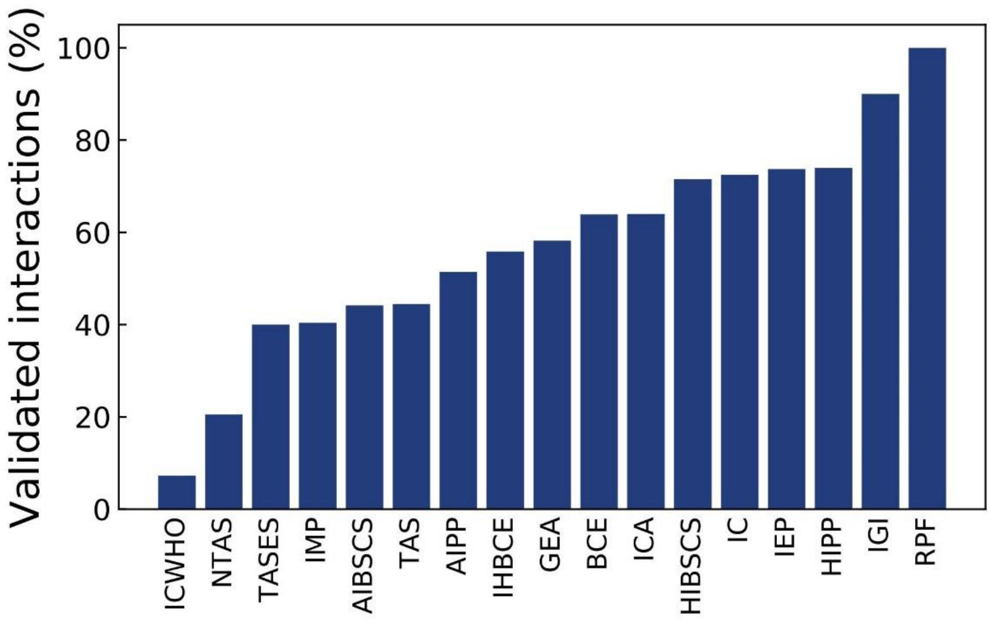


**Supplementary Figure 7.** Interactions inferred by a methodology classified as “weak” that has been validated with a “strong” experiment.

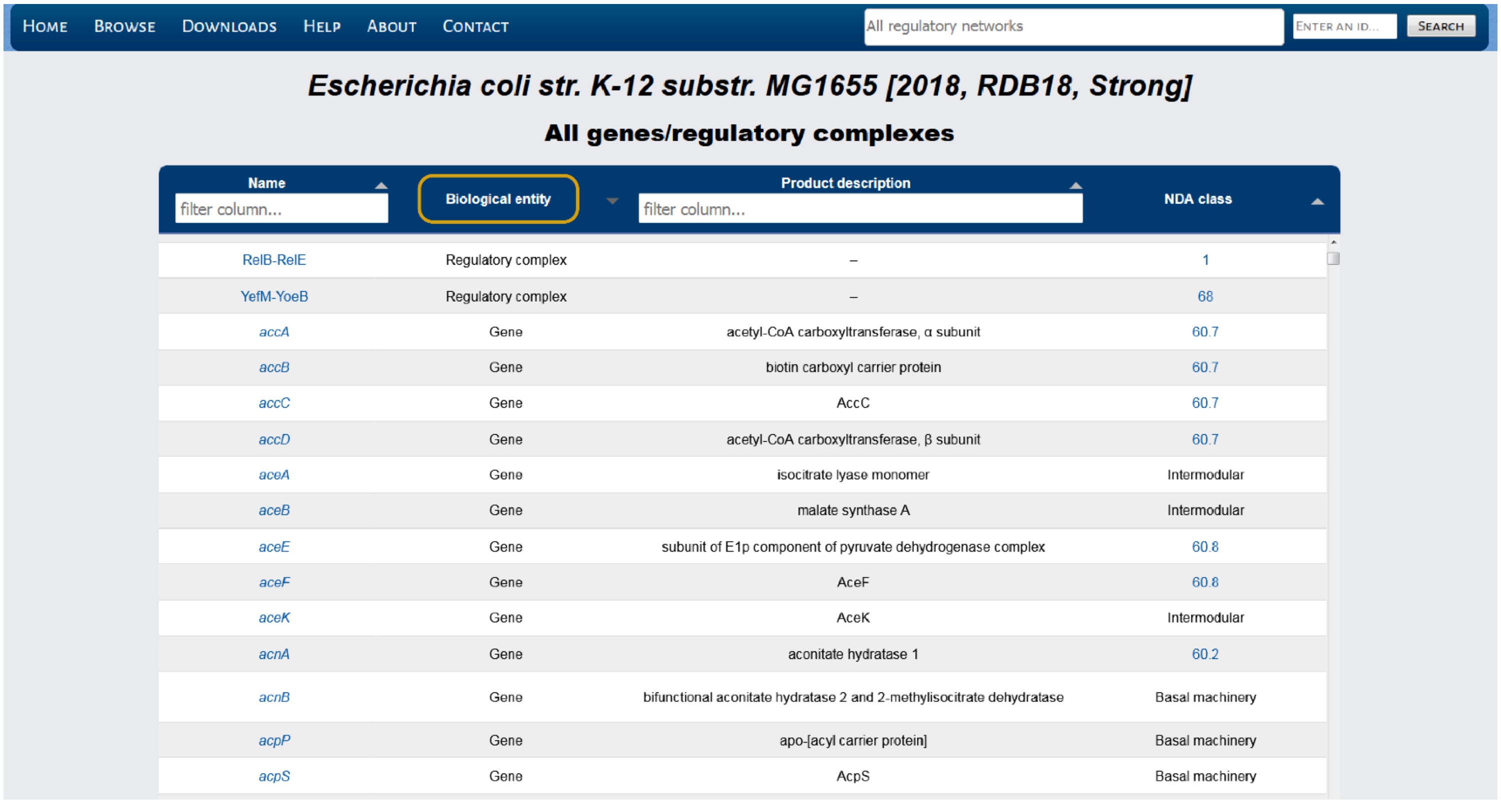


*Escherichia coli str. K-12 substr. MG1655 [2018, RDB18, Strong]*

**All genes/regulatory complexes**

| Name | Biological entity | Product description | NDA class |
|---|---|---|---|
| RelB-RelE | Regulatory complex | – | 1 |
| YefM-YoeB | Regulatory complex | – | 68 |
| *accA* | Gene | acetyl-CoA carboxyltransferase, α subunit | 60.7 |
| *accB* | Gene | biotin carboxyl carrier protein | 60.7 |
| *accC* | Gene | AccC | 60.7 |
| *accD* | Gene | acetyl-CoA carboxyltransferase, β subunit | 60.7 |
| *aceA* | Gene | isocitrate lyase monomer | Intermodular |
| *aceB* | Gene | malate synthase A | Intermodular |
| *aceE* | Gene | subunit of E1p component of pyruvate dehydrogenase complex | 60.8 |
| *aceF* | Gene | AceF | 60.8 |
| *aceK* | Gene | AceK | Intermodular |
| *acnA* | Gene | aconitate hydratase 1 | 60.2 |
| *acnB* | Gene | bifunctional aconitate hydratase 2 and 2-methylisocitrate dehydratase | Basal machinery |
| *acpP* | Gene | apo-[acyl carrier protein] | Basal machinery |
| *acpS* | Gene | AcpS | Basal machinery |

**Supplementary Figure 8.** Regulatory complexes can be identify using the “Biological entity” column and the Gene information file available at the “Downloads” page (see **Supplementary Figure 2)**. This makes possible to convert the GRNs to contain only gene-gene interactions when needed.

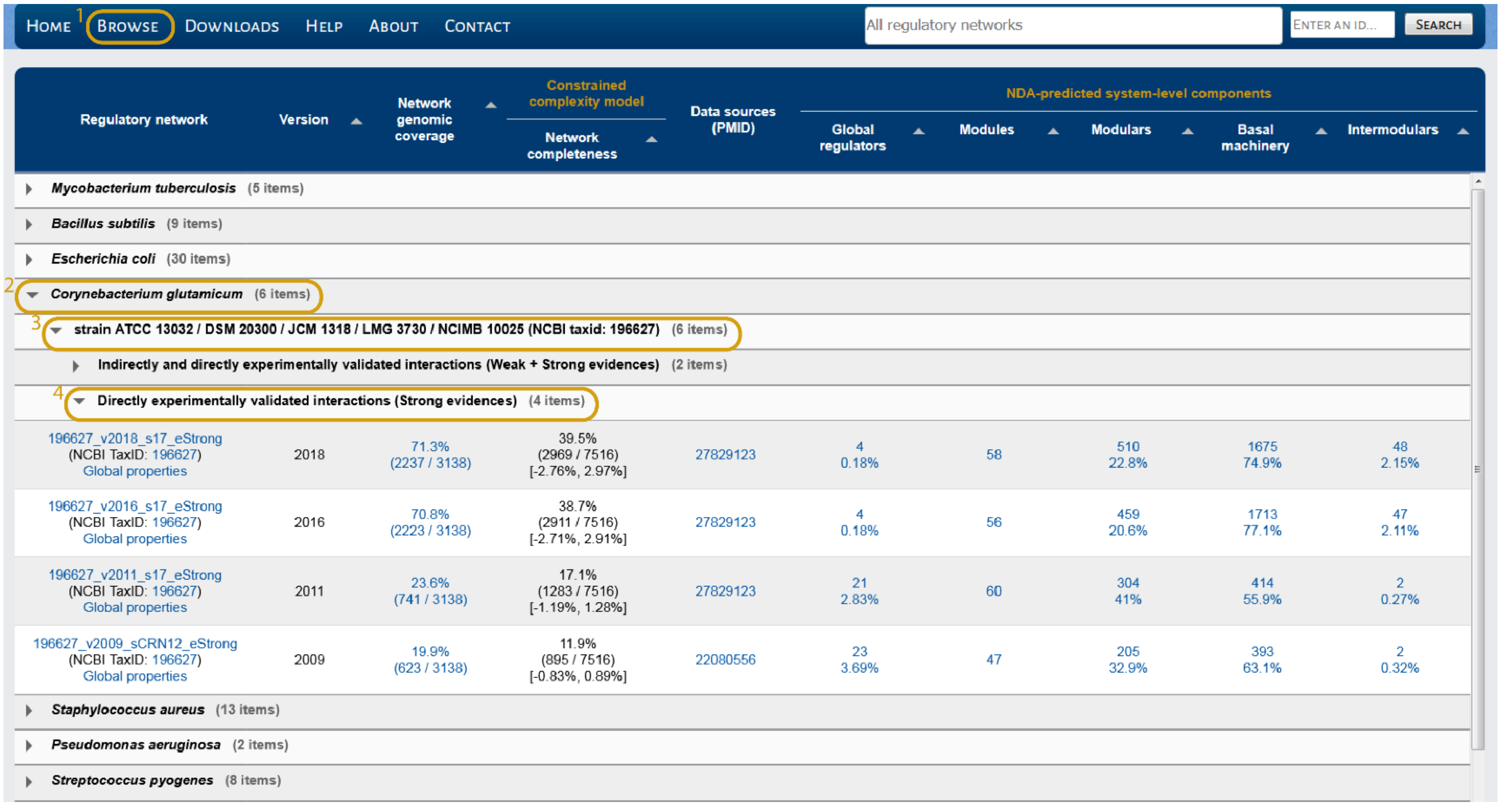


| Regulatory network | Version | Network genomic coverage | Constrained complexity model: Network completeness | Data sources (PMID) | NDA-predicted system-level components: Global regulators | Modules | Modulars | Basal machinery | Intermodulars |
|---|---|---|---|---|---|---|---|---|---|
| ▸ *Mycobacterium tuberculosis* (5 items) | | | | | | | | | |
| ▸ *Bacillus subtilis* (9 items) | | | | | | | | | |
| ▸ *Escherichia coli* (30 items) | | | | | | | | | |
| 2 ▾ *Corynebacterium glutamicum* (6 items) | | | | | | | | | |
| 3 ▾ strain ATCC 13032 / DSM 20300 / JCM 1318 / LMG 3730 / NCIMB 10025 (NCBI taxid: 196627) (6 items) | | | | | | | | | |
| ▸ Indirectly and directly experimentally validated interactions (Weak + Strong evidences) (2 items) | | | | | | | | | |
| 4 ▾ Directly experimentally validated interactions (Strong evidences) (4 items) | | | | | | | | | |
| 196627_v2018_s17_eStrong (NCBI TaxID: 196627) Global properties | 2018 | 71.3% (2237 / 3138) | 39.5% (2969 / 7516) [-2.76%, 2.97%] | 27829123 | 4 0.18% | 58 | 510 22.8% | 1675 74.9% | 48 2.15% |
| 196627_v2016_s17_eStrong (NCBI TaxID: 196627) Global properties | 2016 | 70.8% (2223 / 3138) | 38.7% (2911 / 7516) [-2.71%, 2.91%] | 27829123 | 4 0.18% | 56 | 459 20.6% | 1713 77.1% | 47 2.11% |
| 196627_v2011_s17_eStrong (NCBI TaxID: 196627) Global properties | 2011 | 23.6% (741 / 3138) | 17.1% (1283 / 7516) [-1.19%, 1.28%] | 27829123 | 21 2.83% | 60 | 304 41% | 414 55.9% | 2 0.27% |
| 196627_v2009_sCRN12_eStrong (NCBI TaxID: 196627) Global properties | 2009 | 19.9% (623 / 3138) | 11.9% (895 / 7516) [-0.83%, 0.89%] | 22080556 | 23 3.69% | 47 | 205 32.9% | 393 63.1% | 2 0.32% |
| ▸ *Staphylococcus aureus* (13 items) | | | | | | | | | |
| ▸ *Pseudomonas aeruginosa* (2 items) | | | | | | | | | |
| ▸ *Streptococcus pyogenes* (8 items) | | | | | | | | | |

**Supplementary Figure 9.** From the "Browse" page you can identify the species and strain of interest, as well as the confidence level you need. Also, you will find additional information such as the GC and IC, data sources and fraction of the systems-level components predicted by the NDA. Clicking the hyperlink in the data sources PMID will open the PubMed page of the article for the data source.

HOME BROWSE DOWNLOADS HELP ABOUT CONTACT All regulatory networks ENTER AN ID... SEARCH

### *Mycobacterium tuberculosis (strain ATCC 25618 / H37Rv) [2018, 11-12-15-16, Weak + Strong]*

| Network global properties | | |
|---|---|---|
| Regulators ($k_{out}$ > 0) | – | 179 (5.7%) |
| Structural genes ($k_{out}$ = 0) | – | 2955 (94.3%) |
| Undirected regulatory links | – | 10119 |
| Directed regulatory interactions | – | 10147 |
| Self-regulations | – | 104 (58.1%) |
| Maximum out-connectivity | – | 452 (14.4%) |
| Network density | – | 0.00204 |
| Weakly connected components | – | 3 |
| Genes in the giant component | – | 3128 (99.8%) |
| Feedforward circuits | – | 3994 |
| Complex feedforward circuits | – | 700 |
| 3-Feedback loops | – | 63 |
| Average path length | – | 3.24 |
| Network diameter | – | 6 |
| Average clustering coefficient | – | 0.13008 |
| $P(k)$ | – | $0.772 \cdot k^{-1.97}$ |
| $R^2_{adj}$ | – | 0.93 |

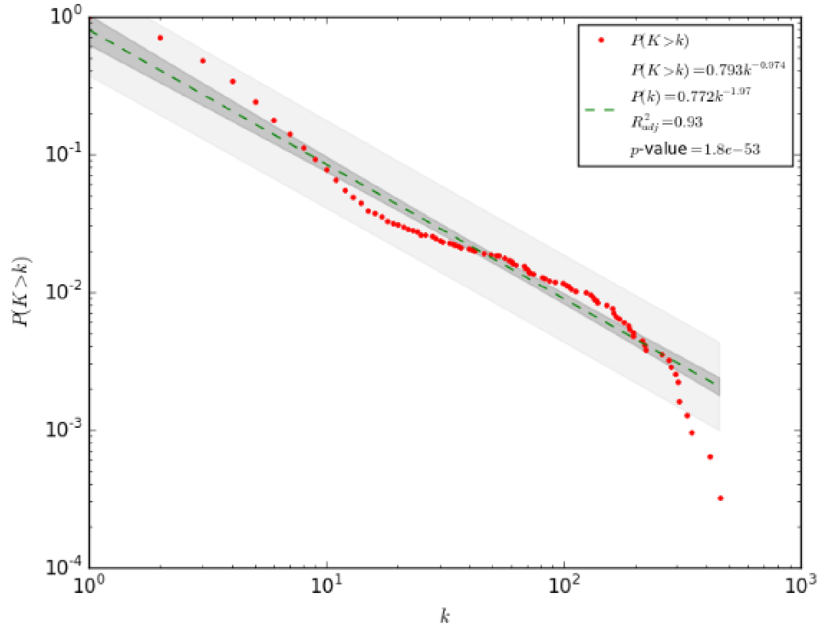


**Supplementary Figure 10.** Statistical and structural properties characterizing the GRN of interest.

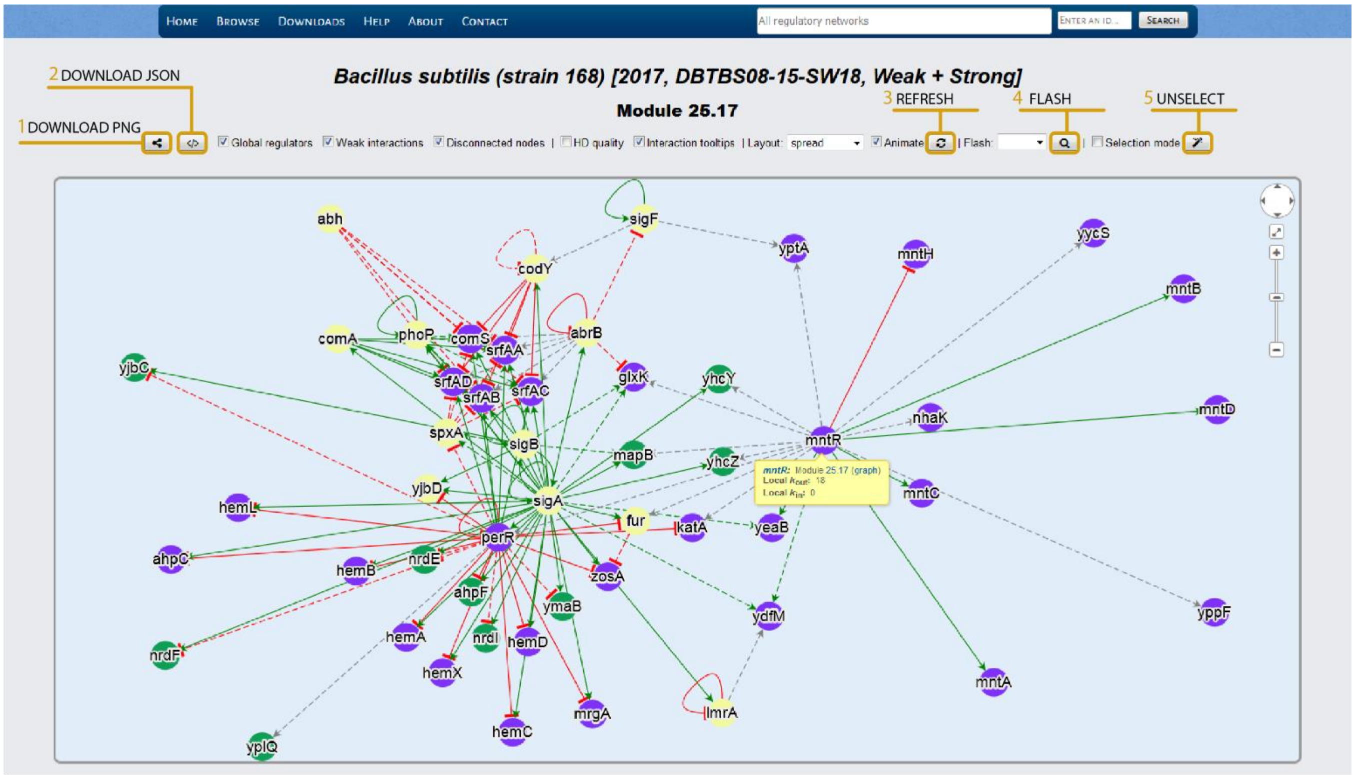


**Supplementary Figure 11.** On the interactive panel you can download the displayed network as an image in PNG format with transparent background (1). Also, you can download the JSON file (2) to import into Cytoscape and customize the style of the network. Check/uncheck the boxes to remove global regulators, "weak" interactions, disconnected genes, interaction tooltips and to display the network in high definition. One the desired boxes are checked, click the refresh button (3) to visualize the changes. Select one gene from the alphabetically sorted "Flash" list and click the flash button (4) to easily identify the gene of interest from the network. Click on the nodes to select them and drag the mouse to change the position of the selected nodes. To deselect all nodes, simply click the unselect button (5).
